# COUNTEN – an AI-driven tool for rapid, and objective structural analyses of the Enteric Nervous System

**DOI:** 10.1101/2020.11.24.396408

**Authors:** Yuta Kobayashi, Alicia Bukowski, Subhamoy Das, Cedric Espenel, Julieta Gomez-Frittelli, Narayani Wagle, Shriya Bakshi, Monalee Saha, Julia A. Kaltschmidt, Archana Venkataraman, Subhash Kulkarni

## Abstract

Healthy gastrointestinal functions require a healthy Enteric Nervous System (ENS). ENS health is often defined by the presence of normal ENS structure. However, we currently lack a comprehensive understanding of normal ENS structure as current methodologies of manual enumeration of neurons within tissue and ganglia can only parse limited tissue regions; and are prone to error, subjective bias, and peer-to-peer discordance. Thus, there is a need to craft objective methods and robust tools to capture and quantify enteric neurons over a large area of tissue and within multiple ganglia. Here, we report on the development of an AI-driven tool COUNTEN which parses HuC/D-immunolabeled adult murine myenteric ileal plexus tissues to enumerate and classify enteric neurons into ganglia in a rapid, robust, and objective manner. COUNTEN matches trained humans in identifying, enumerating and clustering myenteric neurons into ganglia but takes a fraction of the time, thus allowing for accurate and rapid analyses of a large tissue region. Using COUNTEN, we parsed thousands of myenteric neurons and clustered them in hundreds of myenteric ganglia to compute metrics that help define the normal structure of the adult murine ileal myenteric plexus. We have made COUNTEN freely and openly available to all researchers, to facilitate reproducible, robust, and objective measures of ENS structure across mouse models, experiments, and institutions.

---

The myenteric plexus contains the majority of neurons and glial cells of the Enteric Nervous System (ENS) that are clustered in interconnected ganglia of various sizes [1]. The ENS regulates gastrointestinal motility and alterations in ENS structure, gauged by altered neuronal numbers in a defined tissue area or within ganglia, are associated with gastrointestinal dysmotility [2]. While ENS structure is thus relevant to health and disease, methods by which it can be assessed are limited. Currently there are no objective methods to capture and quantify enteric neurons over a large area of tissue or within multiple ganglia. The challenge of objectively quantifying ENS structure at both the neuronal and ganglionic level is particularly acute. While some studies have calculated aggregate myenteric neuronal densities [3-5], they have minimally addressed the ganglionic organization of these neurons (defined here as neuronal numbers per ganglia (ganglia size) along with the density of neurons and ganglia in the tissue). In contrast, recent studies have computed average ganglia size [6, 7], however, due to the laborious process of manually counting and clustering neurons, these studies have analyzed only limited tissue area, thus precluding a comprehensive understanding of structure of the adult myenteric plexus. In addition, since myenteric ganglia show a diversity of shapes and sizes, and exhibit varying degrees of proximity from each other, the current method of manual enumeration and clustering of neurons into ganglia is prone to error, subjective bias, and peer-to-peer discordance. Hence, there is a critical need for objective and rapid methods for standardized enumeration and classification of myenteric neurons into ganglia over large areas to build a comprehensive understanding of ENS structure.

Here, we present the first automated software that can reliably identify and cluster myenteric neurons across a large field of view. Our software, called COUNTEN (COUNTing Enteric Neurons), uses computer-vision and image-processing methods for rapid and high-throughput analysis of widefield microscopy images. COUNTEN algorithm follows a sequence of four steps (Fig 1): (1) image preprocessing using Otsu’s adaptive thresholding method [8] to separate foreground neurons from the background tissue, (2) neuron identification based on peak intensities within a local neighborhood, (3) neuronal clustering into ganglia using the DBSCAN algorithm [9], and (4) image post-processing via watershed segmentation [10]. These four steps provide high-concordance data on the enumeration of neurons present in tissue and within defined ganglia.

**Figure 1:**
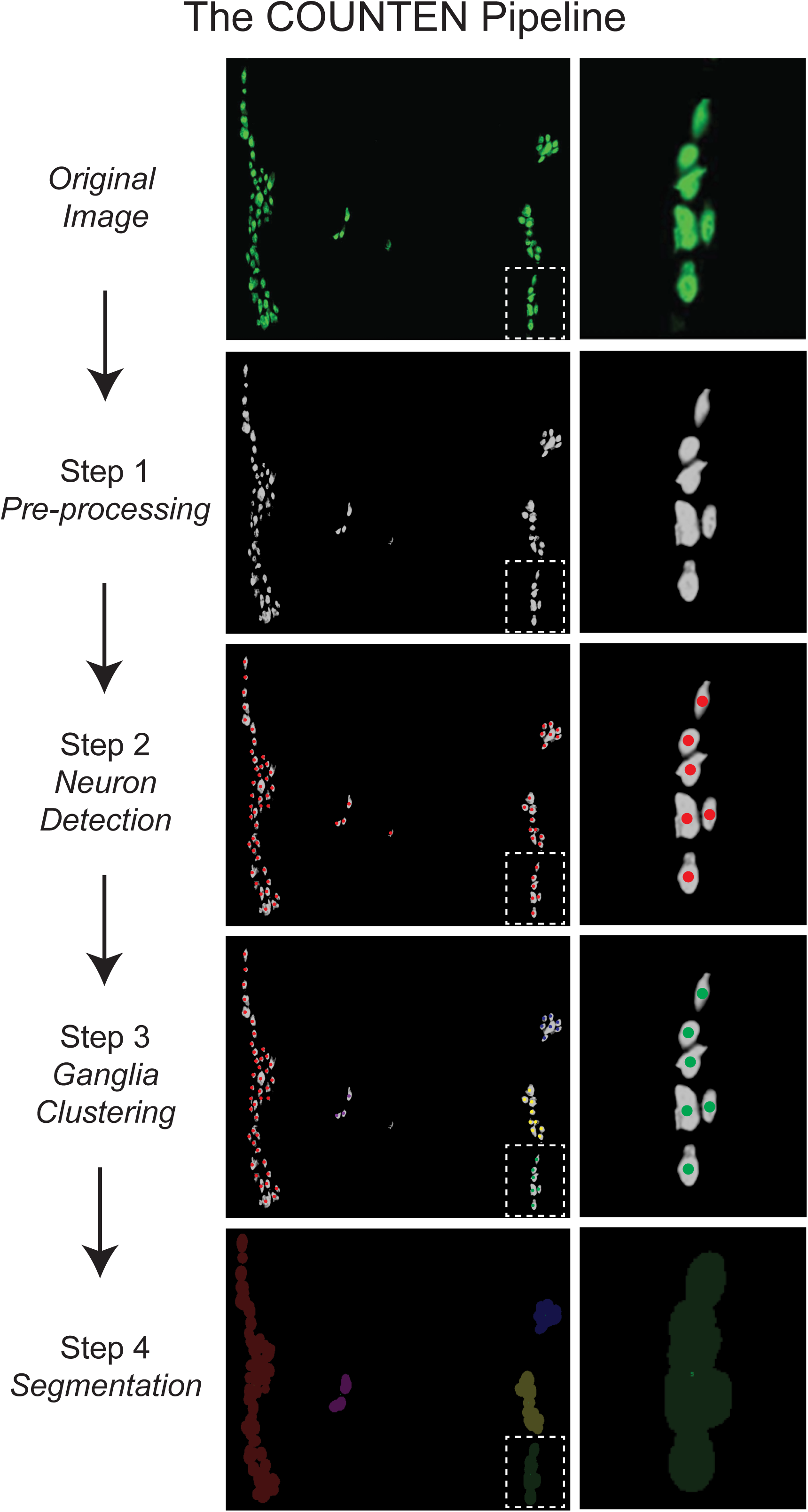
Representative images detailing the automatic COUNTEN image processing sequence for neuronal identification, enumeration, and clustering of HuC/D immunostained iLM-MP in a single 20X field of view. A single ganglion (in dotted box) is expanded on the right to show steps of neuronal identification, enumeration and classification into ganglia.

We used COUNTEN to analyze adult murine ileal Longitudinal Muscle – Myenteric Plexus tissue (henceforth: iLM-MP). We first evaluated whether COUNTEN correctly identified and enumerated neurons with the same precision as human experts. By analyzing 100 images (n = 100) of random 20X fields of view of iLM-MP that was immunostained with antibodies against pan-neuronal marker HuC/D, where each image contained different numbers of neurons, we found that COUNTEN achieves high concordance with manual identification and enumeration of neurons performed by human experts (Fig 2A).

**Figure 2:**
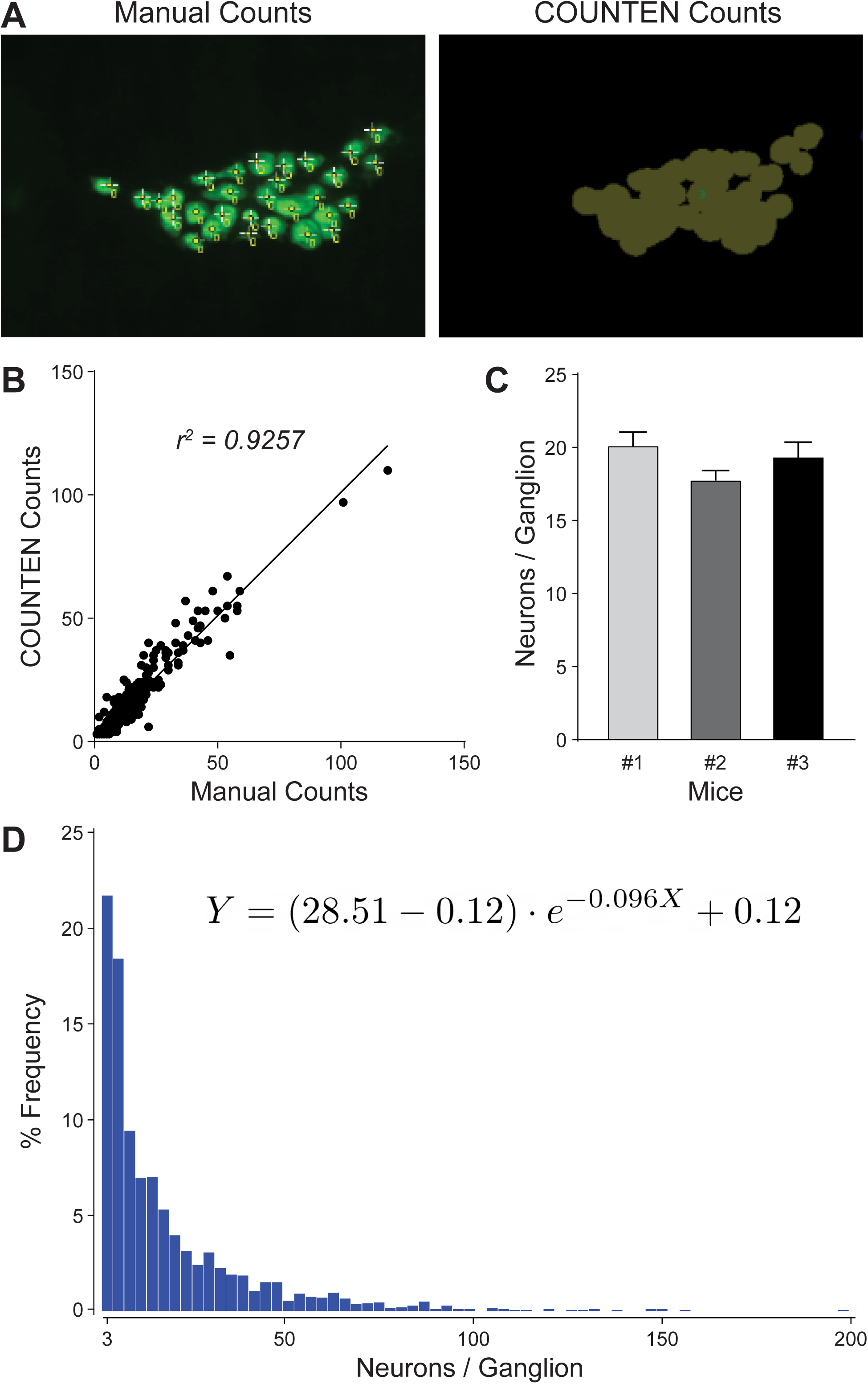
COUNTEN provides rapid, objective, and high-concordance identification, enumeration, and clustering of neurons. (A) Example of human analyzer-driven “Manual” and “COUNTEN”-driven neuronal identification of the same myenteric ganglion. (B) High degree of correlation between COUNTEN-driven enumeration and human analyzer-driven manual enumeration of myenteric neurons per ganglia from the same 100 20X HuC/D immunostained images shows the high degree of conformation between COUNTEN and experienced human analyzer-generated data. (C) COUNTEN-generated data of adult male iLM-MP shows no significant difference in mean numbers of HuC/D-immunostained neurons/ganglia, suggesting a similar ganglia size between litter-mate male mice. Data is represented as Mean ± S.E.M. (D) Frequency distribution histogram of ganglia size shows an inverse correlation between ganglia size and their relative abundance, as represented by the one-phase decay equation. Values on the X axis are in incremental bin sizes of 3 neurons/ganglion.

Second, we evaluated the accuracy of the ganglionic clusters identified by COUNTEN. We defined a ganglion as a cluster of ≥3 neurons. While manual counting identified 413 ganglia in total, COUNTEN achieved similar performance and identified 411 ganglia across the 100 images, underscoring COUNTEN’s reliability. Further, analyses of ganglia size exhibited a similar high degree of concordance between manual and COUNTEN methodologies (*r*^*2*^ = 0.9257; Y=0.9998*X+1.058; Fig 2B). Beyond its reliability, COUNTEN offers a tremendous reduction in the time spent on the analysis. For reference, a trained expert took 2 days to analyze 100 images, while COUNTEN processed the same in <10 minutes, providing us with a platform that performs rapid, precise, and objective neuronal and ganglionic counts in HuC/D-immunostained iLM-MP.

Third, we used COUNTEN to quantify the ganglionic organization of the iLM-MP. We deployed COUNTEN on widefield images generated from multiple stitched 20X images of contiguous fields of view of the HuC/D-immunostained iLM-MP from 3 adult male littermate mice. We imaged an area of 46.15, 48.83, and 36.34 mm^2^ from tissues from the three mice, which were found to contain 15,741, 13,268 and 9,247 neurons within 778, 742, and 475 ganglia, respectively. Using COUNTEN-generated data, we calculated the neuronal density in the three tissues to be 344.33, 288.21, and 269.03 neurons/mm^2^ and the ganglionic density was similarly calculated to be 16.86, 15.19, 13.07 ganglia/mm^2^.

Finally, we used COUNTEN-generated data to study mean ganglia size and their diversity. Ganglia size in neither of the three tissues followed normal distribution (Anderson Darling test, p < 0.0001), and the mean ganglia size (computed to be 20.23, 17.88 and 19.47 neurons/ganglia) between the three tissues were not statistically different (Kruskall-Wallis test) (Fig 2C). Frequency distribution of ganglia size across these three tissues (Fig 2D) showed inverse correlation of ganglia size and their relative abundance, which can be summarized by a one phase decay equation:

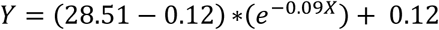

Thus, COUNTEN has provided the first large-scale objective analysis of ileal myenteric neurons and ganglia. The parameters of the one phase decay equation define the frequency distribution of ganglia size, which along with neuronal and ganglionic density provide the metrics of ganglionic organization in the iLM-MP and can be used as a reference for future studies. Biologically, we found that ganglia size is conserved across ileal tissues from sex-matched littermate mice. Importantly, analyses of the same 100 images of immunostained-iLM-MP established a high degree of conformity between human and COUNTEN-generated data (Fig 2B), suggesting that the gains in speed with COUNTEN were not associated with a loss in accuracy with regards to the identification and enumeration of neurons, and their classification into ganglia.

The performance of COUNTEN depends on the robustness and homogeneity of immunostaining and imaging, which we have standardized (see: *Methods*). While ganglia are three-dimensional (3D) structures, and the current algorithm does not operate on 3D image-data, given the high concordance between COUNTEN and human-generated data, the data generated by COUNTEN by parsing standard widefield fluorescence microscopy-based images of myenteric plexus are as accurate as those generated by a human observer.

To facilitate rapid and reproducible measures of ENS structure within the broader ENS community, COUNTEN is freely and openly available to researchers.

## Acknowledgements

We are grateful to the Cell Sciences Imaging Facility (CSIF) at Stanford University for initial program development. This work was supported by a Stanford ChEM-H Seed Grant (S.D.); a Beckman Center Bioimage Analysis grant (C.E.); the Stanford ChEM-H Chemistry/Biology Interface Predoctoral Training Program and the National Institute of General Medical Sciences of the NIH under Award Number T32GM120007 (J.G.-F.); the Wu Tsai Neurosciences Institute and a research grant from The Shurl and Kay Curci Foundation (J.A.K.); grants from the National Science Foundation (CRCNS 1822575 and CAREER award 1845430) (A.V.); and funding from the Ludwig Foundation and a pilot grant from the Hopkins Digestive Diseases Basic & Translational Research Core Center grant P30DK089502 (S.K.).

## Methods

### Animals

All animal experiments were conducted in accordance with the protocols that were approved by the Johns Hopkins University Animal Care and Use Committee in accordance with the guidelines provided by the National Institutes of Health. Nine-week-old littermate male mice from the C57BL/6 (Charles River) background were used for the experiment.

### Tissue isolation

Mice were anesthetized with isoflurane and were sacrificed by cervical dislocation. An abdominal incision was made to perform a laparotomy and isolate intestines that were gently pulled out and placed in a clean petri dish containing sterile ice-cold Opti-MEM solution. The intestinal contents were flushed using ice-cold sterile PBS after which the terminal ileum, defined as the last 5 cm of the small intestinal tissue before cecum, was dissected out. The longitudinal muscle containing myenteric plexus (LM-MP) tissue from the terminal ileum was peeled out with a sterile clean cotton swab, cleaned in sterile ice-cold OptiMEM, flattened on a dish and fixed with freshly prepared ice-cold 4% paraformaldehyde solution for 30 minutes. The tissue was washed in sterile ice-cold PBS and used for immunostaining.

### Immunostaining

Fixed iLM-MP tissues were incubated at room temperature (RT) with shaking in Blocking Permeabilization Buffer (BPB: 5% normal goat serum, 0.1% Triton-X in sterile PBS) after which tissues were washed in sterile PBS and incubated overnight with Rabbit anti-HuC/D primary antibody (1:750; Abcam) at 16°C with constant shaking. The tissues were removed from the primary antibodies, washed thrice (10 minutes each) with PBS and incubated in Goat anti-Rabbit Alexa 488 secondary antibody (Invitrogen) at RT for 1 hour in the dark. Subsequently the tissues were washed thrice (10 minutes each) with PBS and mounted with Prolong Anti-Fade mounting medium containing nuclear stain DAPI (Invitrogen). Care was taken not to let the tissue fold on itself during the mounting process.

### Imaging

Using the EVOS M7000 motorized-stage fluorescent microscope (Thermo-Fisher), the tissues were imaged under a 20X (EVOS AMEP4924; Fluorite LWD, 0.45NA/6.23WD) objective. Imaging was performed such that the entire width of the tissue was imaged over variable length. Care was taken not to image folded tissues. Initial concordance measurements of COUNTEN versus manual counting were done using images of individual fields. Subsequently, individual images were stitched together to generate a composite image that was used for COUNTEN analyses for generating the ENS map.

### Manual Counting

Manual Counting of HuC/D immunostained neurons was performed by a trained technician. Using ImageJ to open the images, the technician used the plugin *Cell Counter* on ImageJ to mark individual HuC/D labeled cells to avoid counting the same cell twice. Classification of neurons into ganglia was done by following the rule of defining a cluster of ≥3 neurons as a ganglion. The total numbers of ganglia per image and the numbers of neurons per ganglia were thus enumerated and tabulated.

### Software

The COUNTEN workflow consists of four sequential steps (Fig. 1): (1) image preprocessing, (2) neuron identification, (3) neuronal clustering into ganglia, and (4) image post-processing for segmentation. This algorithm was implemented in Python using the scikit-image, the NumPy, and the scikit-learn libraries. The same workflow was used for all the processing described in this report. The COUNTEN software is freely available on Github (https://github.com/KLab-JHU/COUNTEN).

COUNTEN requires as input two user-specified parameters: the pixel density *ρ* (pixels/*μm*) as dictated by the imaging protocol, and the full width at half maximum *σ* (pixels) of a Gaussian smoothing kernel used during preprocessing. The four steps of the workflow are detailed below:

1. **Image Preprocessing:** This step eliminates noise and staining variations, which might otherwise confound the results. We opted for a simple procedure, which can easily be replicated across different equipment configurations. The RGB image is first converted to a single grayscale channel and processed using an isotropic Gaussian filter. Larger blurs will reduce the contribution of extra-ganglionic neurons but also make the algorithm more susceptible to false negatives. Smaller blurs may result in insufficient denoising of the image. We have empirically determined that setting the Gaussian full width at half maximum to *σ =* 7 pixels yields highly concordant neuronal counts, as compared to human raters (see Fig. 2). Hence, we fix *σ = 7* for all analyses in this work. Next, we divide the image into nine equal partitions and use the center region to set a threshold between foreground (neurons) and background (GI tract). We use just the center region to avoid biasing the threshold based on abnormalities at the tissue edges. The threshold is selected adaptively using Otsu’s method [8], which minimizes the intra-class variance.
2. **Neuron Identification:** This procedure searches for and returns all local maxima within the image, separated by a distance of at least *δ*_*m*_ (pixels). In other words, the peaks are local maxima of a circular neighborhood in the image with a prespecified radius of *δ*_*m*_. When there are multiple peaks within the same neighborhood, then the average of these coordinates is returned. We have fixed the default value of *δ*_*m*_ according to the pixel density *ρ* as follows:

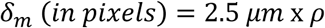 We note that the parameter *δ*_*m*_ is accessible to users within the Python source code and can be modified from this default value as needed for other imaging protocols.
3. **Clustering into Ganglia:** We use the Density-Based Spatial Clustering of Applications with Noise (DBSCAN) algorithm [9] to cluster the peak locations from Step 2 into ganglia. DBSCAN is effective in our application since it does not assume a predefined number of clusters, and it allows for unlabeled points (i.e., extra-ganglionic neurons). The DBSCAN algorithm takes as input two parameters, the minimum number of neurons in a ganglion *N*_*g*_, and the minimum separation between ganglia ε_*m*_. In this work, we fix *N*_*g*_ *= 3* (ganglia contain at least 3 neurons), and this convention was kept constant between COUNTEN-driven neuronal and ganglionic counts and manual counts using human expert. We have fixed the default value of *ε*_*m*_ according to the pixel density *ρ* as follows:

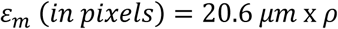 Once again, the parameter *ε*_*m*_ is accessible to users within the Python source code and can be modified from this default setting as needed.
4. **Output Segmentation:** We binarize the image and use the watershed segmentation algorithm [10] to flood the background pixels. This procedure leaves just the identified ganglia as our final output. The algorithm also colors the ganglia for ease of visualization.

